# Codon composition in human oocytes reveals age-associated defects in mRNA decay

**DOI:** 10.1101/2021.01.05.425501

**Authors:** Nehemiah S. Alvarez, Pavla Brachova, Lane K. Christenson

## Abstract

Oocytes from women of advanced reproductive age have lower developmental potential, yet the underlying mechanisms of this phenomena are incompletely understood. Oocyte maturation is dependent upon translational control of stored maternal mRNA that were synthesized during oocyte growth. We observed that GC content of mRNA was negatively associated with half-life in oocytes from reproductively young women (< 30 years), contrastingly directly with oocytes from reproductively aged women (≥ 40 years) where mRNA half-lives were positively associated with GC nucleotide content. Additionally, we observed that mRNA half-lives were negatively associated with protein abundance in young oocytes, while GC content was positively associated with protein abundance in aged oocytes. Examination of codon composition during the GV-to-MII transition revealed that codons that facilitate rapid translation promoted mRNA stability and are considered optimal, while codons that slow translation destabilized mRNA, and are considered non-optimal. GC-containing codons were more optimal in reproductive aging, and also correlated positively with protein abundance. This study indicates that reproductive aging coincides with the stabilization of a subset of mRNA that have the potential to be over-translated during oocyte maturation, this is likely to lead to observed decreases in oocyte quality in older women. Because oocyte mRNA decay is translationally linked, this suggests that maternal aging causes defects in translation, which results in reduced translational efficiency and the retention of maternal mRNA that are normally degraded in oocytes from young women. In the case of oocytes, defects in translation can alter the RNA decay pathways and result in incorrect maternal mRNA dosage, which may negatively impact embryonic development.

## INTRODUCTION

In women, the ovary is the first organ system in the body to display signs of age, a phenomenon known as reproductive aging. At the cellular level, reproductive aging is characterized by a decline in oocyte quality and quantity that results in decreased fertility (Navot et al. 1991). Cellular and molecular mechanisms of human reproductive aging are poorly understood. Aneuploidy is a defining feature associated with reproductive aging in oocytes (Sandalinas et al. 2002; Mai et al. 2013; Treff et al. 2016). Mouse models for reproductive aging have indicated defective translation of meiotic regulators in the oocytes from aged mice, which is hypothesized to drive aneuploidy (Duncan et al. 2017; Llano et al. 2020). The underlying cause of age-associated translational dysregulation remains to be determined in oocytes.

Oocyte growth is characterized by high transcriptional activity, after which mRNA are either translated to synthesize proteins necessary for growth, or stored for future use (Piko and Clegg 1982; Bachvarova et al. 1985; De La Fuente and Eppig 2001). Transcription ceases when oocytes are fully grown, and resumes once the embryonic genome is activated (De La Fuente and Eppig 2001). Thus, oocyte maturation, fertilization, and early embryonic growth rely on stored maternal mRNA. As stored mRNA are recruited for translation, they are subsequently deadenylated, destabilized, and degraded, in a process called translational mRNA decay (Yu et al. 2016; Sha et al. 2018). Therefore, the process of translation ultimately affects mRNA stability. Understanding the intricacies of translational regulation and RNA stability is imperative in order to understand oocyte quality as it relates to reproductive aging.

The sequence of mRNA is known to have important regulatory functions that impact the process of translation (Gruber and Zavolan 2019; Mayya and Duchaine 2019). For example, the 3′ UTR of mRNA has important regulatory capacity in governing the binding of RNA binding proteins and microRNA (Mayya and Duchaine 2019; Harvey et al. 2018). Also, poly(A) tail length plays critical roles in both mRNA stability and translation (Mendez and Richter 2001; Roy and Jacobson 2013). Recently, it has become apparent that the coding sequence (CDS) can also impact important translation-dependent regulatory functions based on codon composition in a phenomenon known as codon optimality (Presnyak et al. 2015; Bazzini et al. 2016; Carneiro et al. 2019; Horstick et al. 2015; Pop et al. 2014; Bergman and Tuller 2020). Enrichment of ‘optimal’ codons tend to stabilize mRNA, display greater abundance, higher translation efficiency, and contain longer poly(A)-tails. Conversely, mRNA enriched in ‘non-optimal’ codons tend to be unstable, have lower abundance, poor translation efficiency, with shorter poly(A)-tails (Bazzini et al. 2016; Mishima and Tomari 2016; Radhakrishnan et al. 2016; Webster et al. 2018). Emerging evidence demonstrates that non-optimal codons can slow translation elongation rates, and decrease mRNA half-life (Nedialkova and Leidel 2015; Presnyak et al. 2015). Specific codons have been associated with mRNA stability (optimal codons) or instability (non-optimal codons) in model organisms, and recently in human cell lines (Presnyak et al. 2015; Boël et al. 2016; Bazzini et al. 2016; Wu et al. 2019; Narula et al. 2019).

The potential for codon composition to regulate translational efficiency and mRNA stability has not yet been examined in human oocytes and in the context of reproductive aging. Here, we assess the coding composition of mRNA during the GV-to-MII transition in human oocytes from reproductively young and aged women to test if the coding sequence impacts mRNA stability and whether translational mRNA decay is impacted by reproductive aging. We observed that during reproductive aging there is a reinforcement of the mRNA decay program within oocytes; with the enhanced stabilization or destabilization of a subset of mRNA. Additionally, we observed that in young oocytes GC-rich codons have a shorter mRNA half-life, and are associated with lower codon optimality and higher protein production in oocytes. During reproductive aging we observed that GC-rich codons have increased codon optimality and GC-rich mRNA have longer half-lives. Thus, codon optimality within oocytes is altered during reproductive aging and this has the potential to alter translational output and oocyte developmental competence.

## MATERIAL AND METHODS

### Alignment and half-life calculation

Reproductively young and aged human GV and MII oocyte RNA-seq data were downloaded from the Short Read Archive project PRJNA377237 (Reyes et al. 2017). These polyA sequencing libraries were generated using SMARTer Ultra Low Input RNA HV kit (Clontech, USA) using the manufacturer’s recommended protocol. Sequences were trimmed of known sequencing adaptors and quality scores using BBMap bbduk with the following settings; maq=30 ktrim=l ktrim=r k=23 mink=11 hdist=1 (Bushnell 2014). Read duplicates with up to 5 mismatches were removed using BBMap clumpify with the following settings; depupe sub=5. Processed reads were aligned to Genecode v34 protein coding transcripts using kallisto v0.45 (Bray et al. 2016). Kallisto data was processed in R for normalization and half-life determination (R Core Team 2013). Alignment statistics are provided in Supplemental File 1.

Transcript per million (TPM) counts were imported into R. Transcripts were eliminated if they had less than 1 TPM and were not reported in every sample for each age group (young GV n=5, young MII n=5, aged GV n=5, aged MII n=5). Filtered data was mean normalized and decay factor corrected as previously described (Sorenson et al. 2018). Standard RNA-seq library normalization strategies such as DESeq and TMM, are inappropriate for RNA decay conditions as both approaches assume between 40-70% of genes are either increasing or decreasing in abundance and the total mass of RNA is relatively constant between conditions (Robinson and Oshlack 2010). In mammalian oocytes, these conditions are not met as GV oocytes are transcriptional quiescent and more than 75% of mRNA is degraded by MII stage (Sha et al. 2018). Decay factor normalization is necessary to account for the decrease in RNA-seq library complexity that occurs in the absence of transcription and the presence of active RNA decay. As a result stable transcripts TPM values will increase in MII oocytes relative to GV oocytes (Sorenson et al. 2018). To perform decay factor normalization, transcript abundance values were normalized to the mean of GV oocytes (time zero) for both reproductively young and aged samples respectively. Next, decay factors were calculated by identifying mRNA whose mean normalized TPM fold increase was greater than one and whose variance across sample groups was less than the mean fold change. The decay factor fold change threshold was empirically determined using an iterative process that incrementally increased the fold change value by 0.5 until the variance of the decay factory normalization reached an inflection point, the fold change value of the inflection point was used as the threshold value for decay factory normalization. Mean normalized and decay factor corrected data were then fit to a linearized version of the following exponential decay model:

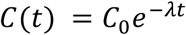

Where, *C*(*t*)is the abundance of RNA at time,*t. C*_*o*_ is the initial abundance of RNA at time zero. *λ*is the decay rate and *t* is time. Decay rates were used to calculate half-life values using the following formula:

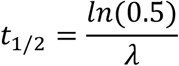

The decay rate, *λ*, was calculated with the following formula:

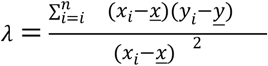

Where, *x*_*i*_, is the time value of the*i*^*th*^ sample, *<u>x</u>*, is the mean of all time, *y*_*i*_, is the abundance value of the*i*^*th*^ sample, *<u>y</u>*, is the mean of sample abundance. Standard error for the decay rate estimation was calculated with the following equation:

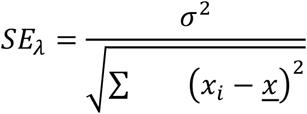

Where *SE*_*λ*_ is the standard error of the estimated decay rate,*x*_*i*_, is the time value of the*i*^*th*^ sample,*<u>x</u>*, is the mean of all time, and σ^2^ is the standard deviation of the variance. The standard deviation of the variance was calculated using the below formula:

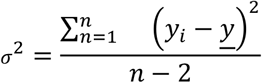

Where, *y*_*i*_, is the abundance value of the*i*^*th*^ sample, *<u>y</u>*, is the mean of sample abundance and*n*, is the degree of freedom. P value estimates for the decay rate were generated by first calculating the t-statistic with the following formula:

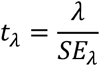

To identify significantly different decay rates between young and aged samples, a standard score was calculated with the following formula:

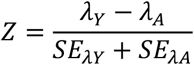

Where, *Z*, is the standard score,*λ*_*Y*_ is the decay rate for a young transcript, *λ*_*A*_ is the decay rate for a aged transcript, *SE*_*λY*_ and *SE*_*λA*_ are the standard error of variance for young and aged mRNA, respectively. The standard score was used to calculate Bonferroni corrected p values.

### mRNA feature, CSC, and pathway analysis

The Genecode v34 protein coding transcript FASTA reference was parsed in R to identify the sequence composition and lengths of CDS and UTR for each transcript. The universal motif Bioconductor package was used to identify GC (nnS, nSS, SSn, SSS) and AU (nnW, nWW, WWn, WWW) containing codons within the CDS of each transcript. The kozak motif (RYMRMVATGGC) was scored in Universalmotif (Noderer et al. 2014). The frequency of codons was calculated for each transcript using the Bioconductor biostrings package. Spearman and Pearson correlations between half-life values and mRNA features were performed in R. The codon correlation coefficient (CSC) was calculated as previously described (Presnyak et al. 2015). The mRNA feature correlations and CSC for reproductive aging samples were calculated by binning the reproductively young half-life values into 1 hour windows. The age half-life values were retrieved for each window, excluding the transcripts that did not increase or decrease more than 1 hour in half-life values. Pathway analysis was performed using ReactomePA in R (Yu and He 2016).

### Mass spectrometry analysis

Mass spectrometry data for human GV and MII oocytes (Virant-Klun et al. 2016) were downloaded from ProteomeXchange Consortium (PXD003691). GV oocyte and MII samples were processed with MaxQuant (v1.6.17.0) and Andromeda search engine as previously described (Virant-Klun et al. 2016; Cox and Mann 2008; Cox et al. 2011). Briefly, the MS/MS spectra were used to search the Human UniProt database (downloaded 07/07/20). Precursor mass tolerance for first pass and second pass were 20 ppm and 6 ppm respectively, while the fragment mass tolerance was set to 0.5 Da and minimum peptide length set to 6. Quantification utilized unmodified unique and razor peptides. Modifications were set to fixed for cysteine carbamidomethylation and methionine oxidation. Trypsin/P was used for enzyme specificity with 2 miss cleavage events. A false discovery rate of 0.01 was used for both peptide and protein identifications. The protein identification was reported as a “protein group” if no unique peptide sequence to a single database entry was identified. Protein abundance was reported as iBAQ values (Schwanhäusser et al. 2011).

## RESULTS AND DISCUSSION

### Age associated changes in mRNA half-life

During the oocyte GV-to-MII transition, changes in mRNA abundance are due to altered mRNA stability rather than mRNA synthesis. To determine the influence of aging on mRNA half-life, we analyzed publicly available human oocyte RNA-seq data from reproductively young and aged women (young: <30 years; aged: ≥ 40 years) (Reyes et al. 2017). Oocyte transcripts accumulate during the growth phase, and are stored in a dormant, untranslated form until meiotic maturation when they become recruited for translation and degradation in a process called translational mRNA decay (Bachvarova et al. 1985; Yu et al. 2016; Sha et al. 2018). We therefore focused our decay analysis on protein coding transcripts that have the potential to be regulated by translational mRNA decay. Previous work has established that a significant number of mammalian transcripts follow an exponential decay pattern (Friedel et al. 2009). Using a mRNA decay normalization strategy, we fit the oocyte polyA RNA sequencing data to an exponential decay curve to calculate mRNA half-life (see methods). We identified 13,421 and 12,794 protein coding transcripts in young and aged oocytes, respectively, undergoing exponential RNA decay (Fig. 1a, p < 0.05, t-statistic, see methods). Of the 11,507 mRNA that overlapped in young and aged oocytes, 6,059 showed a significant difference in half-life (Fig. 1a, p < 0.05, z-score (see methods)). A majority of the protein coding transcripts exhibited a significant increase in half-life during reproductive aging (3,549 increased (Y_1/2_ < A_1/2_); while 2,510 decreased (Y_1/2_ > A_1/2_), Fig. 1b). For young oocyte mRNA that exhibited a decrease in half-life in young vs aged oocytes (Y_1/2_ > A_1/2_), the median half-life was 9.1 hours (Fig 1c, white bar), compared to 9.9 hour median for young oocyte mRNA that exhibited an increase in half-life with aging (Y_1/2_ < A_1/2_; Fig 1c, diagonal bar). This result is somewhat surprising, as one might predict that half-lives for young oocytes in either group (Y_1/2_ < A_1/2_ vs. Y_1/2_ > A_1/2_) would not be significantly different, or that those that were greater in young vs old (Y_1/2_ > A_1/2_) would be more likely to have longer half-lives than those that had reduced half-lives in young vs old (Y_1/2_ < A_1/2_) samples.

**Figure 1.**
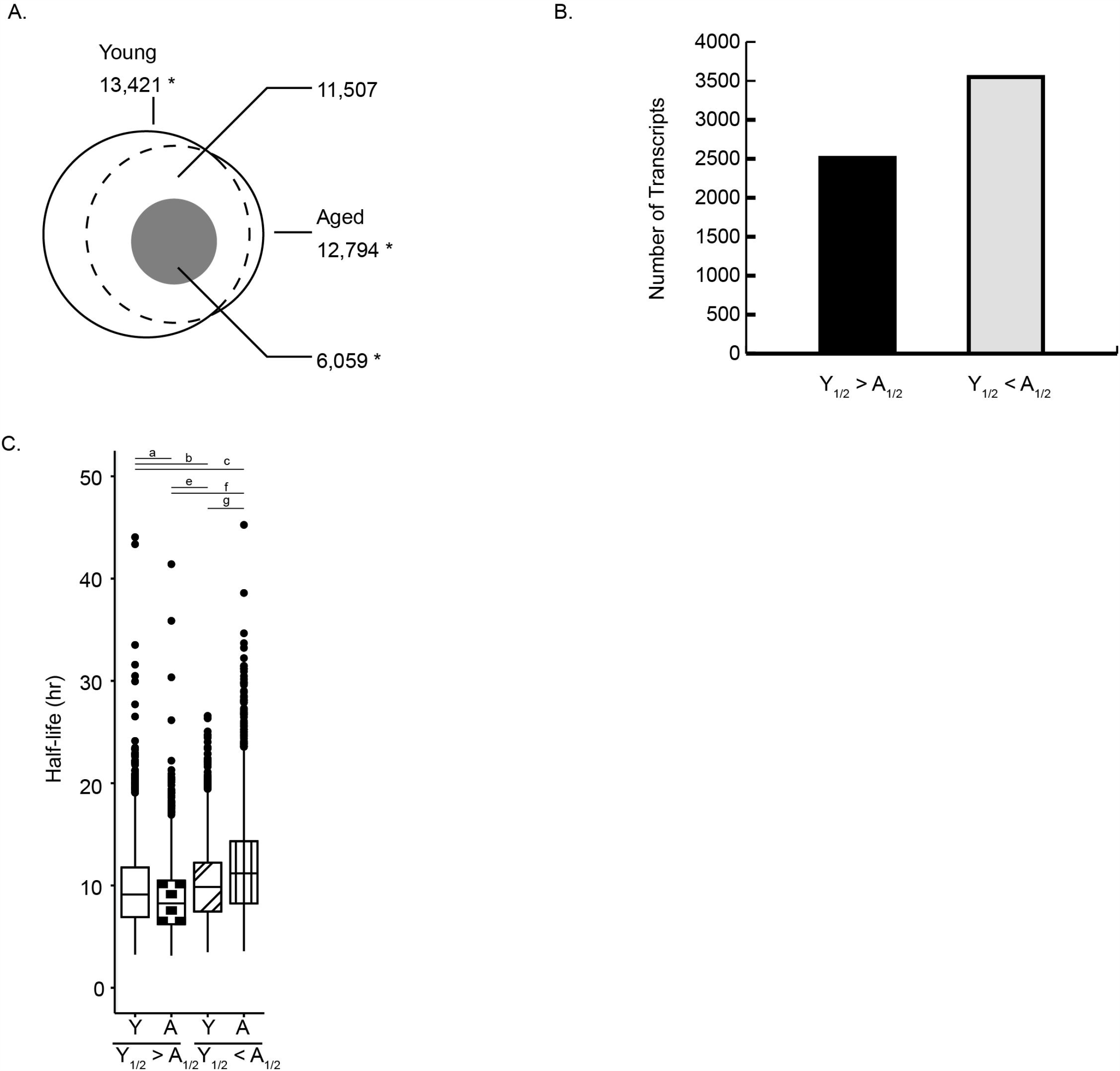
Reproductive aging alters mRNA half-life during oocyte maturation. A) Venn diagram depicts the number of mRNA that fit an exponential decay curve in young and aged oocytes. Dotted circle is the overlap of mRNA between young and aged samples. The gray inset circle represents the common transcripts that have significant differences in half-life between young and aged samples. B) Bar chart depicts the number of mRNA (gray inset from A) with decreased (Y_1/2_ > A_1/2_) or increased (Y_1/2_ < A_1/2_) half-lives associated with reproductive aging. C) Boxplot shows the mRNA half-life for transcripts that either decrease (Y_1/2_ > A_1/2_) or increase (Y_1/2_ < A_1/2_) with reproductive aging. Y: reproductively young, A: reproductively aged. KS test p < 0.05.

Reproductive aging enhances these differences, with aged oocyte mRNA exhibiting an 8.2 hour median half-life (checkered bar) in the Y_1/2_ < A_1/2_ (i.e., short half-life group) and a 11.2 hour median half-life (stripped bar) in the Y_1/2_ > A_1/2_ (i.e., long half-life group). These results suggest that reproductive aging reinforces and enhances mRNA decay programs already present in young oocytes.

### GC content of mRNA positively correlates with half-life during reproductive aging

To gain further insight on the effects of aging on mRNA decay we categorized transcripts into three groups: mRNA that changed less than one hour were deemed to be “not altered” during reproductive aging, or mRNA that either decreased or increased in half-life (Fig. 2a). We observed that a majority of mRNA not altered by aging (white bars) had a half-life of less than 8 hours. Among transcripts with a half-life of longer than 8 hours in reproductively aged samples, the majority were stabilized during reproductive aging, shown as an increased half-life (black bars, Fig. 2a). During human oocyte maturation, the germinal vesicle breaks down at 6-8 hours and oocytes remain arrested at MI for approximately 14 hours (Combelles et al. 2002; Escrich et al. 2012). Theoretically, the increase in half-life of mRNA during aging may impact meiotic competence if particular meiotic transcripts are not degraded at the proper time points. Of the mRNA that increased in half-life during aging, the pathways involved included RNA processing, mitotic processes, microtubule organization from the centrosome, and DNA repair pathways (Supplemental Fig. S1). Of the mRNA that decrease in half-life during aging, those pathways included mitochondrial processes, mitochondrial translation, neddylation and SUMOylation, tRNA processing, and DNA repair (Supplemental Fig. S2). A delay or enhancement of translation and mRNA decay represented in these pathways could negatively impact meiosis and oocyte maturation. Whether the utilization of stored transcript is delayed or hastened, the inappropriate use and decay of maternal mRNA is likely to have profound impacts on oocyte developmental competence.

**Figure 2.**
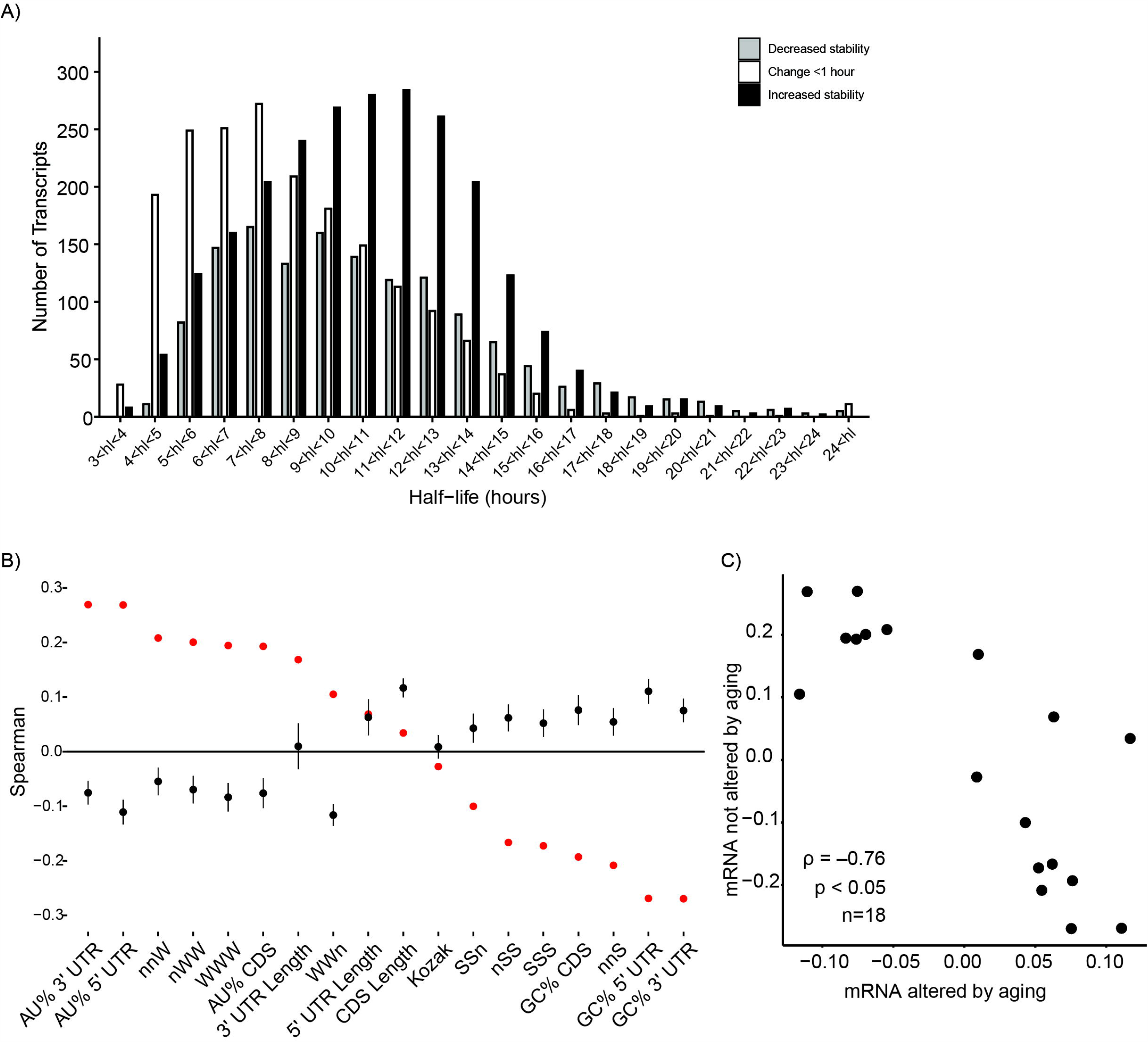
GC content of mRNA positively correlates with half-life during reproductive aging. A) Histogram representing half-life values of decreasing, no change and increasing stability at 1 hour intervals during reproductive aging. B) Depiction of the mean spearman correlation between the mRNA half-life and mRNA features derived from the 1 hour bins in panel A for mRNA not altered by aging (black dots, Mean +/-SEM) and for mRNA altered by aging (red dots, Mean +/-SEM). nSS, nnS, SSS, SSn are IUPAC codons for GC (SS) containing codons. nWW, nWW, WWW, WWn are IUPAC codons for AU (WW) containing codons. C) Spearman correlation between the feature values of B; ρ, Spearman correlation; p, p-value; n, number of features.

Decay of mRNA relies on the sequence composition of its underlying features (Courel et al. 2019). Multiple mRNA features have the potential to impact mRNA decay, including: GC and AU content (in 3’ UTR, 5’ UTR, and CDS) as well as lengths of 3’ UTR, 5’ UTR and CDS, Kozak sequence, and lastly codon composition (Acevedo et al. 2018; Mishima and Tomari 2016). Therefore, we sought to identify features of mRNA that could be associated with alterations to decay during reproductive aging. Global nucleotide composition significantly correlated with mRNA half-life and was impacted by reproductive aging (Fig. 2b). Among mRNA that remained constant during aging, half-life was negatively associated with GC content, while mRNA altered by aging, half-life was positively associated with GC content (Fig. 2b). The half-life/mRNA feature correlation of transcripts not altered in aging has a significant negative relationship with the half-life/mRNA feature correlation of transcripts altered by aging (Fig. 2c, ρ = –0.76, p < 0.05, Spearman). In oocytes, mRNA degradation is coupled to translation and a recent study demonstrated that GC-rich transcripts were highly translated, while AU-rich mRNA were stored (Su et al. 2007; Svoboda et al. 2015; Sha et al. 2019; Courel et al. 2019). For mRNA that exhibit no change in half-life during aging (black dots, Fig. 2b), our results are in agreement with these previous findings showing that AU-rich mRNA have longer half-lives and GC-rich mRNA have shorter half-lives, presumably due to storage versus translational mRNA decay (Courel et al. 2019). However, for mRNA with half-lives affected by reproductive aging, we observe that GC-rich messages have longer half-lives, suggesting a potential defect in translational mRNA decay or mRNA storage occurs with increased age.

### GC content reveals differential correlations between mRNA half-life and protein abundance

To further explore the relationship of mRNA features and translational mRNA decay we compared mRNA half-life changes to oocyte protein abundance using previously generated mass spectrometry data during GV to MII human oocyte maturation (Virant-Klun et al. 2016). We observed an inverse relationship between mRNA half-life and protein abundance (Fig. 3a, ρ = –0.23, p < 0.05). This result indicates that as protein abundance increases, mRNA half-life decreases, which further supports a model of translational mRNA decay in oocytes.

**Figure 3.**
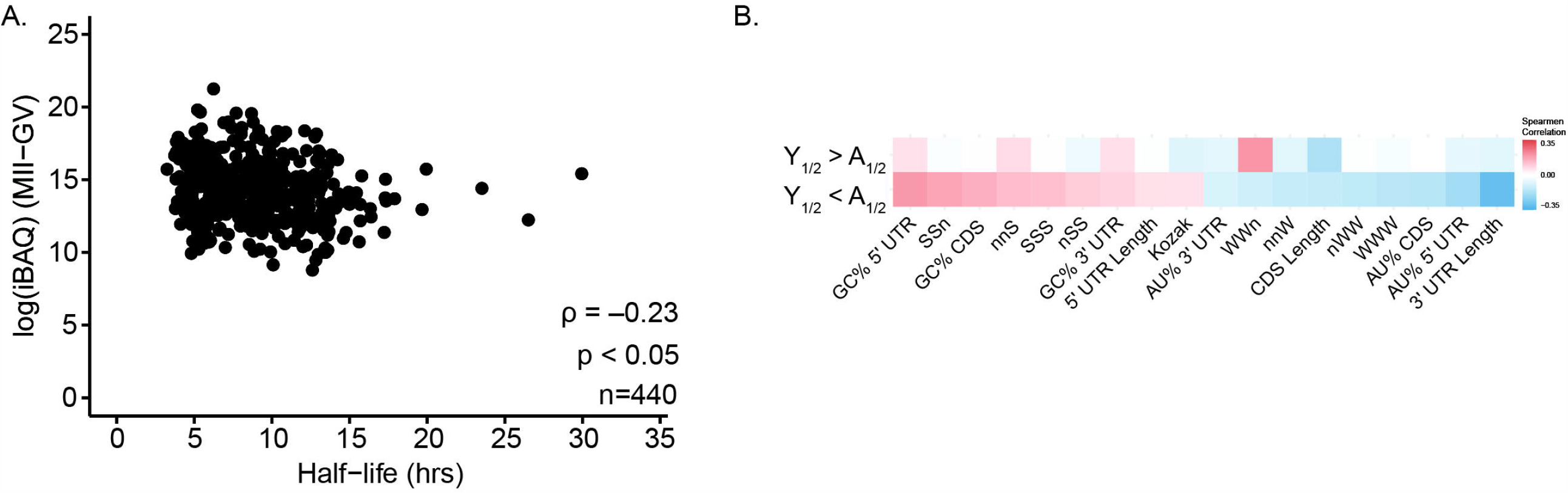
GC/AU content reveals differential correlations between mRNA half-life and protein abundance. A) Scatter plot illustrates relationship between protein abundance (log(iBAQ_MII_-iBAQ_GV_)) and mRNA half-life (reproductively young) from genes (n=440) with a single protein coding isoform; ρ, Spearman correlation; p, p-value. B) Spearman correlation between protein abundance (iBAQ, MII - GV) and mRNA features from corresponding mRNA.

Because reproductive aging enhanced mRNA decay processes already in place (Fig. 1c), we examined mRNA features associated with translation output within mRNA that increase or decrease during reproductive aging. We observed that GC-rich mRNA features within mRNA that increased in half-life during reproductive aging had a positive association with protein abundance (Fig. 3b). Conversely, mRNA that exhibited a decrease in half-life during reproductive aging had a reduced association with GC-content (Fig. 3b). Therefore, transcripts that are stabilized during reproductive aging are GC-rich, and are associated with high protein output. Whether or not these transcripts are similarly translated during reproductive aging remains to be determined. Recent work in mice has shown that reproductive aging in oocytes is associated with increased ribosome synthesis (Duncan et al. 2017) and altered mRNA translation (Llano et al. 2020). It is conceivable that human reproductive aging follows similar aging-related dysregulation, in which there is enhancement of the established mRNA decay programs.

### Age associated changes in codon optimality

Our analysis revealed that GC-containing codons (nSS, nnS, SSS, SSn) were positively associated with protein abundance (Fig. 3) and increased mRNA half-life during reproductive aging (Fig. 1). Experimental evidence indicates that codon composition of an mRNA can influence translation rates, which in turn influence mRNA stability (Presnyak et al. 2015; Bazzini et al. 2016; Carneiro et al. 2019; Horstick et al. 2015; Pop et al. 2014; Bergman and Tuller 2020). This phenomenon termed codon optimality is an emerging contributor to the regulation of mRNA stability in mammals (Bazzini et al. 2016; Wu et al. 2019; Narula et al. 2019). In order to examine codon optimality in oocytes, we calculated the defined metric, codon stability coefficient (CSC; (Presnyak et al. 2015), for mRNA that were affected or unaffected by reproductive aging (Fig. 4a & b). After filtering out neutral codons (0.1> CSC > –0.1; (Carneiro et al. 2019), we observed that for mRNA unaffected by aging, optimal codons were AU enriched: 19 out of 20 codons (Fig. 4a), in contrast to mRNA affected by aging, optimal codons were GC enriched: 24 out of 29 codons (Fig. 4b). Thus, the CSC values for transcripts affected by reproductive aging were negatively correlated with the CSC from mRNA unaffected by aging (Fig. 4c, ρ = –0.62, p < 0.05). For mRNA that is not affected by aging, our codon optimality results are consistent with previous reports in mammalian oocytes and cell lines showing AU-rich codons are associated with mRNA storage (Su et al. 2007; Svoboda et al. 2015; Sha et al. 2019; Courel et al. 2019). For mRNA affected by reproductive aging GC-rich codons are more optimal, indicating potential defects in mRNA storage or translational mRNA decay.

**Figure 4.**
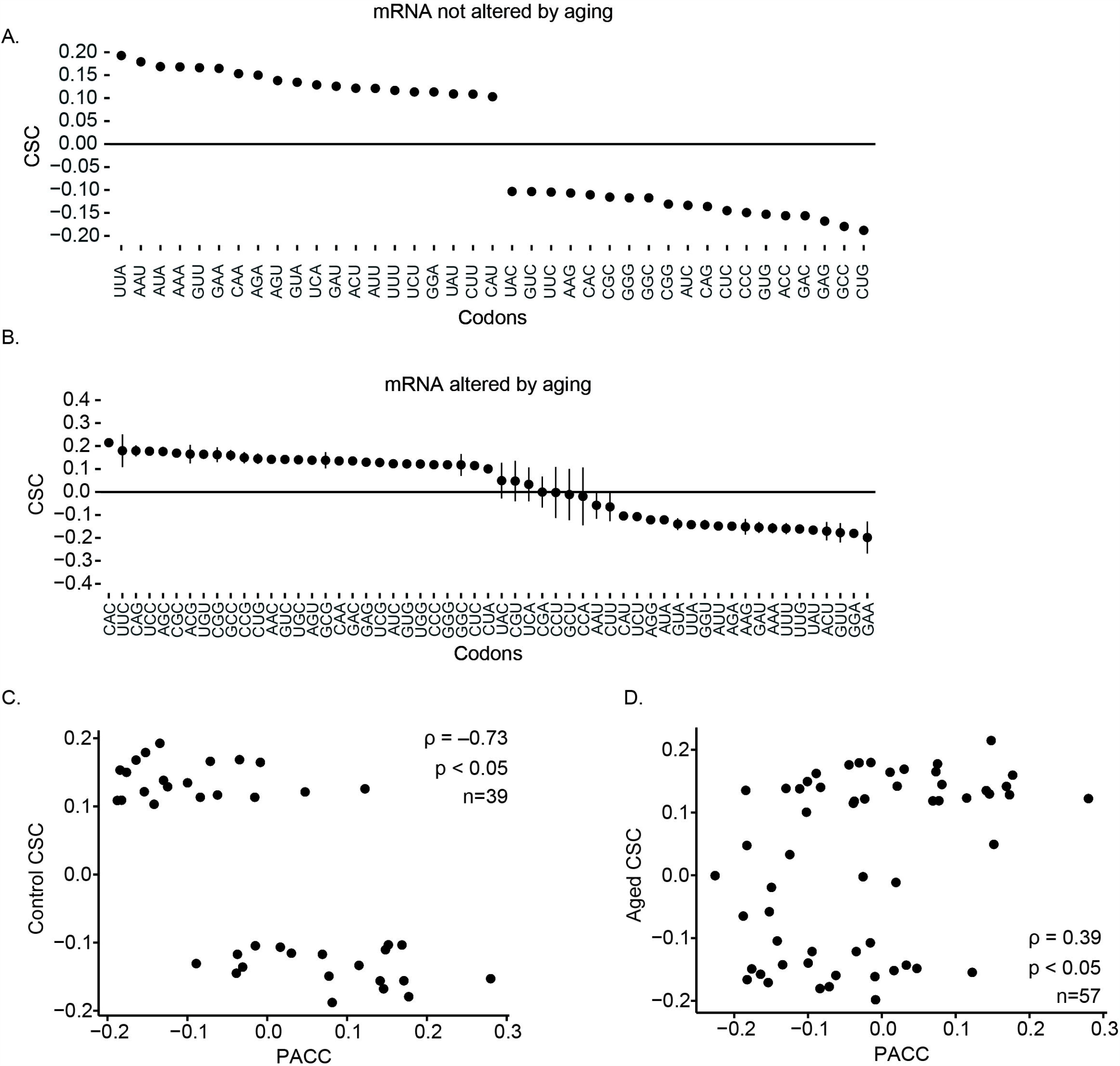
Reproductive aging in human oocytes alters codon optimality. A) CSC from mRNA not altered by reproductive aging. The CSC is an average of CSC values from each 1 hour bin from Fig 2a. CSC values whose value was between −0.1 and 0.1 were filtered out, prior to averaging the CSC, error bars represent the SEM. B) CSC from mRNA altered by reproductive aging. CSC is an average of CSC values from each 1 hour bin from Fig 2a. CSC values whose value was between −0.1 and 0.1 were filtered out, prior to averaging the CSC, error bars represent the SEM. C) Scatter plot comparing the CSC from unaffected mRNA (panel A) to protein abundance codon correlation (PACC); ρ, Spearman correlation; p, p-value; n, number of codons in common. D) Scatter plot comparing the CSC from affected mRNA (panel B) to protein abundance codon correlation (PACC); ρ, Spearman correlation; p, p-value; n, number of codons in common.

In model organisms, the current model explaining the relationship between mRNA stability and codon usage is that some codons (low CSC score and non-optimal) are translated slowly, triggering mRNA decay. We thus examined the relationship of CSC and protein abundance, a relationship we termed the *protein abundance codon correlation* (PACC). The PACC is calculated by substituting the iBAQ (iBAQ_MII_ - iBAQ_GV_) for the mRNA half-life value in the CSC calculation. We compared the CSC of mRNA unaffected by aging to the PACC and observed a negative correlation (Fig. 4c, ρ = –0.73, p < 0.05). On the other hand, the PACC had a positive relationship to CSC derived from mRNA affected by reproductive aging (Fig. 4d, ρ = 0.39, p < 0.05).The correlation between the PACC and the reproductive aging CSC suggest that a subset of mRNA could be potentially stabilized and translated over a longer period of time during oocyte maturation. This phenomenon is called over-translation and has been observed in mice models for defects in translational mRNA decay (Sha et al. 2018) as well as reproductively aged oocytes (Llano et al. 2020).

## Conclusion

Age-associated declines in female fertility are attributed to the combined effects of reduced gamete quantity and quality (Navot et al. 1991). Clinical data from assisted reproductive technology cycles have shown that when women use their own eggs to conceive, the likelihood of having life offspring decreases with age (Check et al. 2011). When women use eggs from young donors, their fertility is restored to that of young women. These studies indicate that the major determinant of fertility outcomes is the biological age of the egg, rather than the woman. Molecular mechanisms responsible for decreased egg quality are multifactorial, and include alterations in chromosome structure, chromosome-associated proteins, cell cycle regulation, and the spindle machinery, and mitochondrial function (Hunt and Hassold 2008; Jones and Lane 2013; Hornick et al. 2015; Seidler and Moley 2015). A recent study in growing mouse follicles identified altered proteostasis and protein processing in advanced age (Duncan et al. 2017). Here, we add evidence to the dysregulation of proteostasis during reproductive aging caused by changes in translational mRNA decay. We observed that in reproductive aging, mRNA GC content was associated with longer half-life and increased translational output. Although the RNA-seq data used in our analysis was from IVF patients whose GV oocytes failed to mature following hCG administration and were then rescued with in vitro maturation (Reyes et al. 2017; Virant-Klun et al. 2016), rescue in vitro maturation can result in pregnancies and live births (Lee et al. 2016; Elanchezhian et al. 2018).In the future it will be highly valuable to assess human reproductive aging in oocytes from healthy individuals in order to identify molecular signatures of advanced maternal age.

## Acknowledgements

This work was supported by the National Institutes of Health (HD094545 to L.K.C., HD099269 to P.B., and HD094545-01A1S1 to N.S.A.). The authors also acknowledge the University of Kansas Medical Center Internal Support and Genomics Core and their funding: Kansas Intellectual and Developmental Disabilities Research Center (NIH U54 HD090216).

## References

Acevedo JM, Hoermann B, Schlimbach T, Teleman AA. 2018. Changes in global translation elongation or initiation rates shape the proteome via the Kozak sequence. Sci Rep 8: 4018.

Bachvarova R, De Leon V, Johnson A, Kaplan G, Paynton BV. 1985. Changes in total RNA, polyadenylated RNA, and actin mRNA during meiotic maturation of mouse oocytes. Dev Biol 108: 325–331.

Bazzini AA, Del Viso F, Moreno-Mateos MA, Johnstone TG, Vejnar CE, Qin Y, Yao J, Khokha MK, Giraldez AJ. 2016. Codon identity regulates mRNA stability and translation efficiency during the maternal-to-zygotic transition. EMBO J 35: 2087–2103.

Bergman S, Tuller T. 2020. Widespread non-modular overlapping codes in the coding regions. Phys Biol. http://dx.doi.org/10.1088/1478-3975/ab7083.

Boël G, Letso R, Neely H, Price WN, Wong K-H, Su M, Luff J, Valecha M, Everett JK, Acton TB, et al. 2016. Codon influence on protein expression in E. coli correlates with mRNA levels. Nature 529: 358–363.

Bray NL, Pimentel H, Melsted P, Pachter L. 2016. Near-optimal probabilistic RNA-seq quantification. Nat Biotechnol 34: 525–527.

Bushnell B. 2014. BBMap: a fast, accurate, splice-aware aligner. Lawrence Berkeley National Lab.(LBNL), Berkeley, CA (United States) https://www.osti.gov/biblio/1241166.

Carneiro RL, Requião RD, Rossetto S, Domitrovic T, Palhano FL. 2019. Codon stabilization coefficient as a metric to gain insights into mRNA stability and codon bias and their relationships with translation. Nucleic Acids Res 47: 2216–2228.

Check JH, Jamison T, Check D, Choe JK, Brasile D, Cohen R. 2011. Live delivery and implantation rates of donor oocyte recipients in their late forties are similar to younger recipients. J Reprod Med 56: 149–152.

Combelles CMH, Cekleniak NA, Racowsky C, Albertini DF. 2002. Assessment of nuclear and cytoplasmic maturation in in-vitro matured human oocytes. Hum Reprod 17: 1006– 1016.

Courel M, Clément Y, Bossevain C, Foretek D, Cruchez OV, Yi Z, Bénard M, Benassy M-N, Kress M, Vindry C, et al. 2019. GC content shapes mRNA storage and decay in human cells. eLife 8. http://dx.doi.org/10.7554/elife.49708.

Cox J, Mann M. 2008. MaxQuant enables high peptide identification rates, individualized p.p.b.-range mass accuracies and proteome-wide protein quantification. Nat Biotechnol 26: 1367–1372.

Cox J, Neuhauser N, Michalski A, Scheltema RA, Olsen JV, Mann M. 2011. Andromeda: a peptide search engine integrated into the MaxQuant environment. J Proteome Res 10: 1794–1805.

De La Fuente R, Eppig JJ. 2001. Transcriptional activity of the mouse oocyte genome: companion granulosa cells modulate transcription and chromatin remodeling. Dev Biol 229: 224–236.

Duncan FE, Jasti S, Paulson A, Kelsh JM, Fegley B, Gerton JL. 2017. Age-associated dysregulation of protein metabolism in the mammalian oocyte. Aging Cell 16: 1381– 1393.

Elanchezhian M, Sankari S, Selvamani D, Nagarajan M, Gopikrishnan D. 2018. Live birth after rescue in vitro maturation–intracytoplasmic sperm injection in type 1 diabetes, polycystic ovary syndrome patient using clomiphene–antagonist protocol. Journal of Human Reproductive Sciences 11: 75. http://dx.doi.org/10.4103/jhrs.jhrs_65_17.

Escrich L, Grau N, de los Santos MJ, Romero J-L, Pellicer A, Escribá M-J. 2012. The dynamics of in vitro maturation of germinal vesicle oocytes. Fertil Steril 98: 1147–1151.

Friedel CC, Dölken L, Ruzsics Z, Koszinowski UH, Zimmer R. 2009. Conserved principles of mammalian transcriptional regulation revealed by RNA half-life. Nucleic Acids Res 37: e115.

Gruber AJ, Zavolan M. 2019. Alternative cleavage and polyadenylation in health and disease. Nat Rev Genet 20: 599–614.

Harvey RF, Smith TS, Mulroney T, Queiroz RML, Pizzinga M, Dezi V, Villenueva E, Ramakrishna M, Lilley KS, Willis AE. 2018. Trans-acting translational regulatory RNA binding proteins. Wiley Interdiscip Rev RNA 9: e1465.

Hornick JE, Duncan FE, Sun M, Kawamura R, Marko JF, Woodruff TK. 2015. Age-associated alterations in the micromechanical properties of chromosomes in the mammalian egg. J Assist Reprod Genet 32: 765–769.

Horstick EJ, Jordan DC, Bergeron SA, Tabor KM, Serpe M, Feldman B, Burgess HA. 2015. Increased functional protein expression using nucleotide sequence features enriched in highly expressed genes in zebrafish. Nucleic Acids Res 43: e48.

Hunt PA, Hassold TJ. 2008. Human female meiosis: what makes a good egg go bad? Trends Genet 24: 86–93.

Jones KT, Lane SIR. 2013. Molecular causes of aneuploidy in mammalian eggs. Development 140: 3719–3730.

Lee H-J, Barad DH, Kushnir VA, Shohat-Tal A, Lazzaroni-Tealdi E, Wu Y-G, Gleicher N. 2016. Rescue in vitro maturation (IVM) of immature oocytes in stimulated cycles in women with low functional ovarian reserve (LFOR). Endocrine 52: 165–171.

Llano E, Masek T, Gahurova L, Pospisek M, Koncicka M, Jindrova A, Jansova D, Iyyappan R, Roucova K, Bruce AW, et al. 2020. Age-related differences in the translational landscape of mammalian oocytes. Aging Cell 19: 1547.

Mai CT, Kucik JE, Isenburg J, Feldkamp ML, Marengo LK, Bugenske EM, Thorpe PG, Jackson JM, Correa A, Rickard R, et al. 2013. Selected birth defects data from population-based birth defects surveillance programs in the United States, 2006 to 2010: featuring trisomy conditions. Birth Defects Res A Clin Mol Teratol 97: 709–725.

Mayya VK, Duchaine TF. 2019. Ciphers and Executioners: How 3’-Untranslated Regions Determine the Fate of Messenger RNAs. Front Genet 10: 6.

Mendez R, Richter JD. 2001. Translational control by CPEB: a means to the end. Nat Rev Mol Cell Biol 2: 521–529.

Mishima Y, Tomari Y. 2016. Codon Usage and 3’ UTR Length Determine Maternal mRNA Stability in Zebrafish. Mol Cell 61: 874–885.

Narula A, Ellis J, Taliaferro JM, Rissland OS. 2019. Coding regions affect mRNA stability in human cells. RNA 25: 1751–1764.

Navot D, Bergh PA, Williams MA, Garrisi GJ, Guzman I, Sandler B, Grunfeld L. 1991. Poor oocyte quality rather than implantation failure as a cause of age-related decline in female fertility. Lancet 337: 1375–1377.

Nedialkova DD, Leidel SA. 2015. Optimization of Codon Translation Rates via tRNA Modifications Maintains Proteome Integrity. Cell 161: 1606–1618.

Noderer WL, Flockhart RJ, Bhaduri A, Diaz de Arce AJ, Zhang J, Khavari PA, Wang CL. 2014. Quantitative analysis of mammalian translation initiation sites by FACS-seq. Mol Syst Biol 10: 748.

Piko L, Clegg KB. 1982. Quantitative changes in total RNA, total poly(A), and ribosomes in early mouse embryos. Dev Biol 89: 362–378.

Pop C, Rouskin S, Ingolia NT, Han L, Phizicky EM, Weissman JS, Koller D. 2014. Causal signals between codon bias, mRNA structure, and the efficiency of translation and elongation. Mol Syst Biol 10: 770.

Presnyak V, Alhusaini N, Chen Y-H, Martin S, Morris N, Kline N, Olson S, Weinberg D, Baker KE, Graveley BR, et al. 2015. Codon optimality is a major determinant of mRNA stability. Cell 160: 1111–1124.

Radhakrishnan A, Chen Y-H, Martin S, Alhusaini N, Green R, Coller J. 2016. The DEAD-Box Protein Dhh1p Couples mRNA Decay and Translation by Monitoring Codon Optimality. Cell 167: 122–132.e9.

R Core Team. 2013. R: A language and environment for statistical computing. http://www.R-project.org/.

Reyes JM, Silva E, Chitwood JL, Schoolcraft WB, Krisher RL, Ross PJ. 2017. Differing molecular response of young and advanced maternal age human oocytes to IVM. Hum Reprod 32: 2199–2208.

Robinson MD, Oshlack A. 2010. A scaling normalization method for differential expression analysis of RNA-seq data. Genome Biol 11: R25.

Roy B, Jacobson A. 2013. The intimate relationships of mRNA decay and translation. Trends Genet 29: 691–699.

Sandalinas M, Márquez C, Munné S. 2002. Spectral karyotyping of fresh, non-inseminated oocytes. Mol Hum Reprod 8: 580–585.

Schwanhäusser B, Busse D, Li N, Dittmar G, Schuchhardt J, Wolf J, Chen W, Selbach M. 2011. Global quantification of mammalian gene expression control. Nature 473: 337– 342.

Seidler EA, Moley KH. 2015. Metabolic Determinants of Mitochondrial Function in Oocytes. Semin Reprod Med 33: 396–400.

Sha Q-Q, Yu J-L, Guo J-X, Dai X-X, Jiang J-C, Zhang Y-L, Yu C, Ji S-Y, Jiang Y, Zhang S-Y, et al. 2018. CNOT6L couples the selective degradation of maternal transcripts to meiotic cell cycle progression in mouse oocyte. EMBO J. http://dx.doi.org/10.15252/embj.201899333.

Sha Q-Q, Zhang J, Fan H-Y. 2019. A story of birth and death: mRNA translation and clearance at the onset of Maternal-to-Zygotic transition in mammals. Biol Reprod. http://dx.doi.org/10.1093/biolre/ioz012.

Sorenson RS, Deshotel MJ, Johnson K, Adler FR, Sieburth LE. 2018. Arabidopsis mRNA decay landscape arises from specialized RNA decay substrates, decapping-mediated feedback, and redundancy. Proc Natl Acad Sci U S A 115: E1485–E1494.

Su Y-Q, Sugiura K, Woo Y, Wigglesworth K, Kamdar S, Affourtit J, Eppig JJ. 2007. Selective degradation of transcripts during meiotic maturation of mouse oocytes. Dev Biol 302: 104–117.

Svoboda P, Franke V, Schultz RM. 2015. Sculpting the Transcriptome During the Oocyte-to-Embryo Transition in Mouse. Curr Top Dev Biol 113: 305–349.

Treff NR, Krisher RL, Tao X, Garnsey H, Bohrer C, Silva E, Landis J, Taylor D, Scott RT, Woodruff TK, et al. 2016. Next Generation Sequencing-Based Comprehensive Chromosome Screening in Mouse Polar Bodies, Oocytes, and Embryos. Biol Reprod 94: 76.

Virant-Klun I, Leicht S, Hughes C, Krijgsveld J. 2016. Identification of Maturation-Specific Proteins by Single-Cell Proteomics of Human Oocytes. Mol Cell Proteomics 15: 2616– 2627.

Webster MW, Chen Y-H, Stowell JAW, Alhusaini N, Sweet T, Graveley BR, Coller J, Passmore LA. 2018. mRNA Deadenylation Is Coupled to Translation Rates by the Differential Activities of Ccr4-Not Nucleases. Mol Cell 70: 1089–1100.e8.

Wu Q, Medina SG, Kushawah G, DeVore ML, Castellano LA, Hand JM, Wright M, Bazzini AA. 2019. Translation affects mRNA stability in a codon-dependent manner in human cells. Elife 8. http://dx.doi.org/10.7554/eLife.45396.

Yu C, Ji S-Y, Sha Q-Q, Dang Y, Zhou J-J, Zhang Y-L, Liu Y, Wang Z-W, Hu B, Sun Q-Y, et al. 2016. BTG4 is a meiotic cell cycle-coupled maternal-zygotic-transition licensing factor in oocytes. Nat Struct Mol Biol 23: 387–394.

Yu G, He Q-Y. 2016. ReactomePA: an R/Bioconductor package for reactome pathway analysis and visualization. Mol Biosyst 12: 477–479.

